# Exercise training improves exercise capacity independent of AMPKα2 T172-mediated adaptations in skeletal muscle

**DOI:** 10.64898/2026.06.18.733224

**Authors:** Xuansong Mao, Ryan N. Montalvo, Kenya Takahashi, Frank W. Booth, George A. Brooks, Zhen Yan

## Abstract

Regular exercise induces adaptations in skeletal muscle and other organ systems to improve physical performance and overall health. Exercise results in phosphorylation of 5’ AMP-activated protein kinase (AMPK) at threonine 172 (T172) of the α2 subunit; however, the role of this activation in cellular and functional adaptations has not been elucidated. To this end, we subjected non-activatable *Ampkα2*(*T172A*) knock-in (KI) adult mice and wild-type (WT) littermates to 4 weeks of voluntary wheel running (VWR). Exercise training led to significant improvements in endurance capacity, maximal oxygen consumption (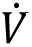O_2_max), and glucose tolerance, as well as skeletal muscle IIb-to-IIa fiber type shift in both WT and KI mice. Contrastingly, VWR resulted in increased mitochondrial OxPhos protein expression, mitochondrial volume density, and capillary density in skeletal muscle of WT but not KI mice. Exercise-induced improvements of mitochondrial respiration and conductance revealed by high-resolution respirometry of isolated mitochondria were blunted in KI mice. Therefore, for the first time, we reveal that AMPKα2 T172 activation is required for exercise training-induced mitochondrial biogenesis, improvement of mitochondrial respiratory function, and angiogenesis in skeletal muscle, but that these adaptations are not solely responsible for improved 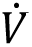O_2_max and exercise endurance capacity.

**Significance Statement:** Exercise is the most effective lifestyle intervention for promoting health and preventing chronic diseases through adaptive changes in skeletal muscle and many other tissues/organs. AMPK is an energy sensor and signaling regulator for exercise-induced skeletal muscle adaptation, yet its functional role and the impact on exercise capacity have been studied in mouse genetic models wherein protein stoichiometry is disrupted. Using non-activatable *Ampkα2(T172A)* knock-in mice, we ascertained that AMPKα2 activation via T172 phosphorylation is required for endurance training-induced mitochondrial and angiogenic adaptations in skeletal muscle. Importantly, these adaptations are not required for improved exercise capacity, challenging the prevailing concept that increased mitochondrial content and function and microvasculature are the sole driving factors for the performance gains with endurance training.

## Introduction

A fundamental theory of exercise physiology is that the human body adapts to the acute and chronic stress of physical activity to improve its maximal functional capacity (1). Exercise training, particularly endurance exercise, leads to adaptations in most systems, including the cardiovascular and musculoskeletal systems that support an overall increase in exercise capacity and performance (2, 3). Important clinically, exercise capacity, or cardiorespiratory fitness, is inversely associated with all-cause of mortality (4). Exercise training is now considered the most potent lifestyle intervention for health promotion and chronic disease prevention (5–7).

Extensive research has been directed toward skeletal muscle adaptations responsible for improved exercise capacity and metabolic homeostasis. In a seminal study, John Holloszy (1967) demonstrated that in rodents, long-term treadmill running can double mitochondrial enzyme activities in skeletal muscle, providing the first biochemical evidence of exercise-induced mitochondrial adaptation (8). The results of increased skeletal muscle mitochondrial content and electron transport chain enzymatic activity have since been reaffirmed in a vast number of studies (2, 9–11), thus laying the foundation for the idea that mitochondrial expansion is crucial for improved exercise endurance capacity after training (12–14). In another seminal study at the same time period, Carrow et al. (1967) showed that long-term treadmill running in rats increased capillary density in skeletal muscle (15), which has since been reaffirmed (16, 17). Intriguingly, these two lines of concurrent research focused on cellular structures that are responsible for the delivery and consumption of oxygen as energy system essential for ATP production to main function of skeletal muscle. Since then, significant focus has been given to the mass of the mitochondrial oxidative energy system and capillary remodeling in skeletal muscle (18–28). The findings have led to a long-standing concept that mitochondrial and microvascular adaptations in skeletal muscle are crucial for improved exercise capacity and performance induced by exercise training (12, 13).

In the last three decades, numerous studies have focused on identification and elucidation of signaling regulators for mitochondrial biogenesis and angiogenesis in skeletal muscle in response to exercise training. 5’ AMP-activated protein kinase (AMPK), an energy sensor and regulator of metabolism (29–34), emerged as a primary determinant of skeletal muscle adaptation. AMPK is readily activated by phosphorylation at threonine 172 (T172) of the dominant α2 catalytic subunit in response muscle contraction and exercise (35, 36). In support of this conclusion, muscle-specific overexpression of a dominant-negative form of Ampkα2 resulted in diminished exercise-induced increases in mitochondrial content and mitochondria-specific antioxidant capacity (37). Muscle-specific deletion of a AMPK downstream target gene, peroxisome-proliferator-activated receptor γ coactivator 1α (*Pgc-1α*) (38), blocked exercise training-induced mitochondrial biogenesis and angiogenesis in skeletal muscle (39, 40); whereas muscle-specific overexpression of Pgc-1α promoted mitochondrial biogenesis and angiogenesis in skeletal muscle (41, 42). Furthermore, skeletal muscle-specific deletion of the upstream AMPK kinase, Lkb1, blunted exercise training-induced mitochondrial biogenesis in skeletal muscle (43). In contrast, there are findings that do not support the inferred importance of AMPK in skeletal muscle adaptation. For example, mice with muscle-specific deletion of the *Ampkα2* gene or the *Ampkα1*/*Ampkα2* genes, or muscle-specific overexpression of an inactive Ampkα2 showed normal enhancement of mitochondrial biogenesis after training (44–47). Hence, the definitive roles of AMPK activation in skeletal muscle adaptation to exercise training remains to be fully elucidated.

The interpretations of some of these previous findings may have been influenced by the methodology of gene deletion or dominant-negative overexpression approaches, which can alter protein stoichiometry (i.e., loss or overexpression of specific proteins) that have been shown to cause compensatory changes (48–50). Furthermore, prior works using these genetic models with exercise training have not comprehensively measured mitochondrial respiratory capacity, metabolic flexibility and free radical generation, which are key determinants of mitochondrial OXPHOS coupling efficiency, respiratory control, and overall energy substrate partitioning.

To evaluate the role of AMPKα2 T172 activation in skeletal muscle adaptations in response to endurance training, we employed *Ampkα2(T172A)* knock-in (KI) mice generated through CRISPR-Cas9-mediated gene editing by substituting the T172 phosphorylation for alanine (T172A) in the Ampkα2 subunit (51). We hypothesized that AMPKα2 T172 activation in skeletal muscle is required for exercise-induced mitochondrial biogenesis and angiogenesis, which are necessary for the improved endurance exercise capacity and VO_2_max. For this reason, we comprehensively examined the role of AMPKα2 T172 activation in skeletal muscle mitochondrial adaptation at protein, structural and functional levels. We also assessed angiogenesis and fiber-type shift and determined how these adaptive processes regulated by AMPKα2 T172 activation contribute to the exercise performance.

## Results

### *Ampkα2(T172A)* adult KI mice exhibit normal whole-body functional adaptations in response to endurance exercise training

Exercise-induced AMPK activation in skeletal muscle has been well documented by studies in animals and humans (36, 52). Consistent with this, we observed robust AMPKα1/2 activation through T172 phosphorylation in plantaris muscle by low-frequency (10 Hz, 2 hr) electric stimulation of the plantar flexor muscles (Fig. S1*A-B*), as well as 90-min treadmill running (Fig. S1*C*-*I*), along with activation of downstream signaling molecules (Fig. S1*A*-*I*). As described above, to ascertain the role of AMPKα2 T172 activation in exercise training-induced adaptations, we uniquely employed non-activatable *Ampkα2(T172A)* KI mice in this experiment (Fig. 1*A, S2A-B*). Homozygous KI and their wild type (WT) littermates were subjected to 4 weeks of voluntary wheel running (VWR). On average, KI mice exhibited reduced voluntary running distance compared to WT mice (WT: 10.4 vs. KI: 7.4 km/day, Fig. 1*B*, S2*C*).

**Figure 1.**
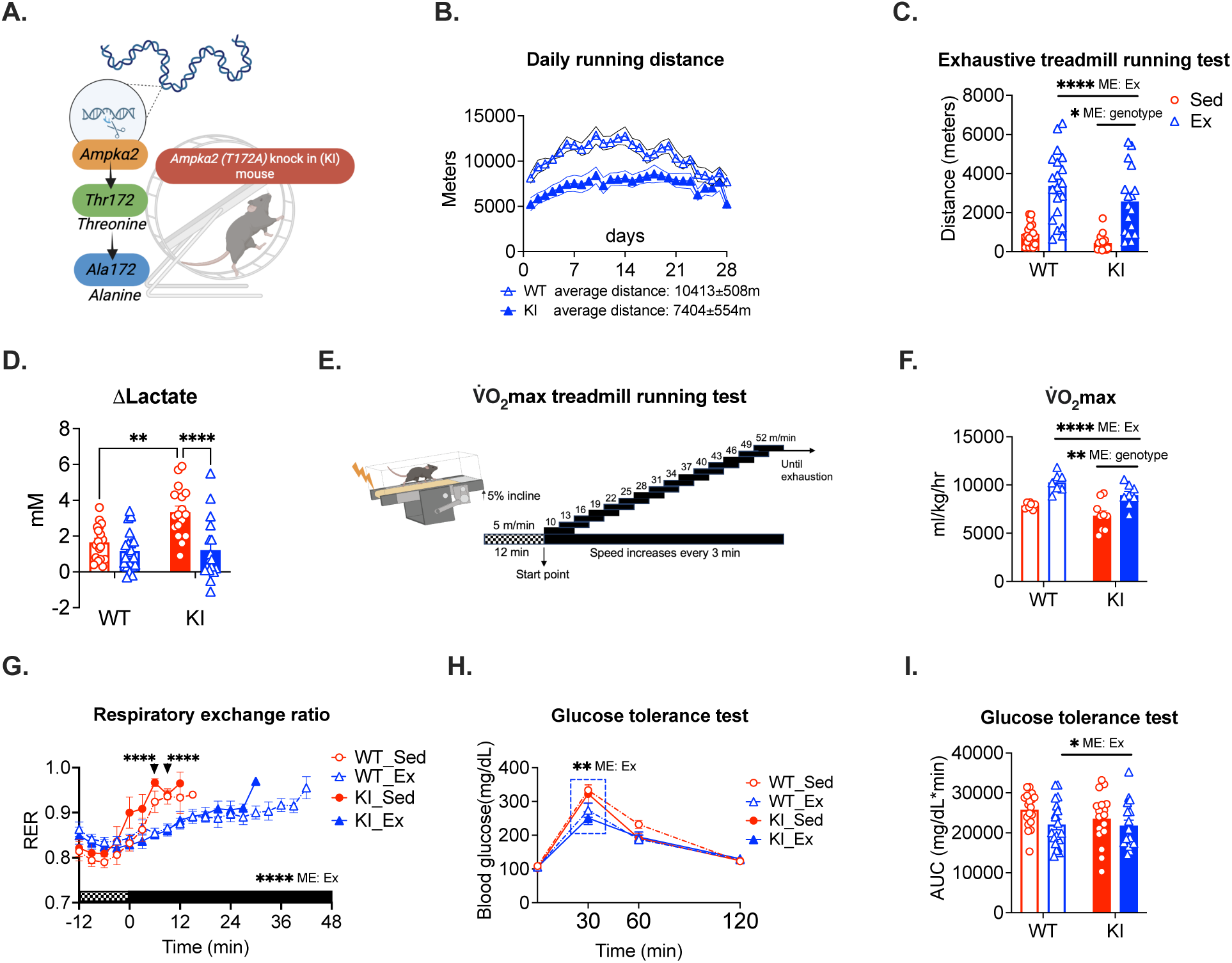
Exercise training improves exercise capacity and glucose tolerance independent of Ampkα2 T172 activation. *Ampkα2*(T172A) KI and WT littermates were subjected to VWR for 4 weeks with sedentary KI and WT mice as controls followed by exhaustive treadmill running test, VO_2_max treadmill running test and GTT. **A**: Schematic illustration of *Ampkα2*(T172A) KI mouse; **B**: Daily voluntary wheel running distance for WT (10413±508m) and KI (7404±554m) mice in exercise groups; **C**: Running distance of exhaustive treadmill running test for all groups; **D:** Changes of blood lactate before and after exhaustive treadmill running test, n = 18-20 per group**; E**: Graphic demonstration of VO_2_max treadmill running test; **F**: Maximal oxygen consumption (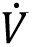O_2_max); **G**: Respiratory exchange ratio (RER) (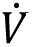CO_2_/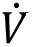O_2_), n = 7-9 per group; **H**: Blood glucose during GTT; and **I**: Area under the curve during GTT in all groups, n = 18-20 per group. Data presented as means ± SEM analyzed by two-way ANOVA. \**p* < 0.05, \*\**p* < 0.01, \*\*\*\**p* < 0.0001. ME: main effect; Ex: exercise.

We assessed exercise capacity by an exhaustive treadmill running test (ETT). As shown recently (51), current sedentary KI mice had impaired running capacity compared to WT controls (Fig. 1*C*). VWR improved running distance significantly in both WT and KI mice (Fig. 1*C*). Elevated blood lactate at the end of ETT confirmed exhaustion in both animal groups (Fig. S2*D*). The significantly greater blood lactate post-ETT, shown as Δlactate in sedentary KI mice, is interpreted to mean greater lactate production via glycogenolysis and glycolysis and/or limited lactate clearance capacity (Fig. 1*D*) (53, 54), To further evaluate the impact on exercise capacity and metabolism, we performed a VO_2_max treadmill running test (Fig. 1*E*) to assess maximal oxygen consumption (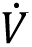O_2_max), energy substrate utilization partitioning via respiratory exchange ratio (RER) (i.e., VCO_2_/VO_2_). VWR significantly improved maximal running performance, 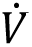O_2_max, maximum speed, heat production, but not maximal RER, in both WT and KI mice compared to sedentary controls (Fig. 1*F*, S2*E-I*). VWR also delayed the rise of RER in both KI and WT mice (Fig. 1*G*), indicating increased utilization on fatty acid metabolism during submaximal exercise intensities (Fig. S2*J* and *K*). VWR improved glucose tolerance in both WT and KI groups, reflected by both reduced area under the curve and reduced elevation of blood glucose at 30 min after glucose injection (Fig. 1*H* and *I*). In addition, VWR reduced body weight and fat mass without affecting lean mass as well as moderately decreased gastrocnemius muscle mass and increased soleus muscle mass in both WT and KI mice, (Fig. S3*A*-*H*). Collectively, despite reduced voluntary running distance, *Ampkα2(T172A)* KI mice had normal improvements in endurance capacity and metabolic adaptations following endurance training.

### AMPKα2 T172 activation is required for exercise-induced mitochondrial adaptations and angiogenesis but not for fiber-type transformation in skeletal muscle

Given the exercise performance and metabolic findings, we next examined parameters of mitochondrial biogenesis, fiber type transformation and angiogenesis, which are key adaptations in skeletal muscle following endurance training (8, 10, 55–58). Protein markers of mitochondrial content, including cytochrome c oxidase subunit 4 (Cox4) and cytochrome c (Cyt c), were significantly downregulated in *Ampkα2(T172A)* KI mice at baseline, while VWR led to a trend of increased expression in WT mice but not in KI mice (Fig. 2*A*-*C*). Similar but less robust trends were observed for mitochondrial membrane voltage-dependent anion channel protein (Vdac), and mitochondria-specific antioxidant enzyme, superoxide dismutase 2 (Sod2) (Fig. 2*D*-*E*). Mitochondrial OXPHOS subunit proteins followed similar trends, with complex IV protein significantly elevated in WT mice but not in KI mice after VWR (Fig. 2*F* and *G*). Our findings suggest the requirement of AMPKα2 T172 activation in enhanced expression of key mitochondrial respiratory components by exercise. To assess proteins involving in mitochondrial structural organization, we examined components of the mitochondrial contact site and cristae organizing system (MICOS) (59). Core components of Mic60-containing subcomplex (Immt and Apool) were reduced KI mice, whereas Mic19 (Chchd3) and Samm50, a component of the sorting and assembly machinery (SAM) complex were unchanged (Fig. S4*A-E*). Our findings suggest that disrupted cristae organization may also underlie impaired mitochondrial function in the skeletal muscle of KI mice.

**Figure 2.**
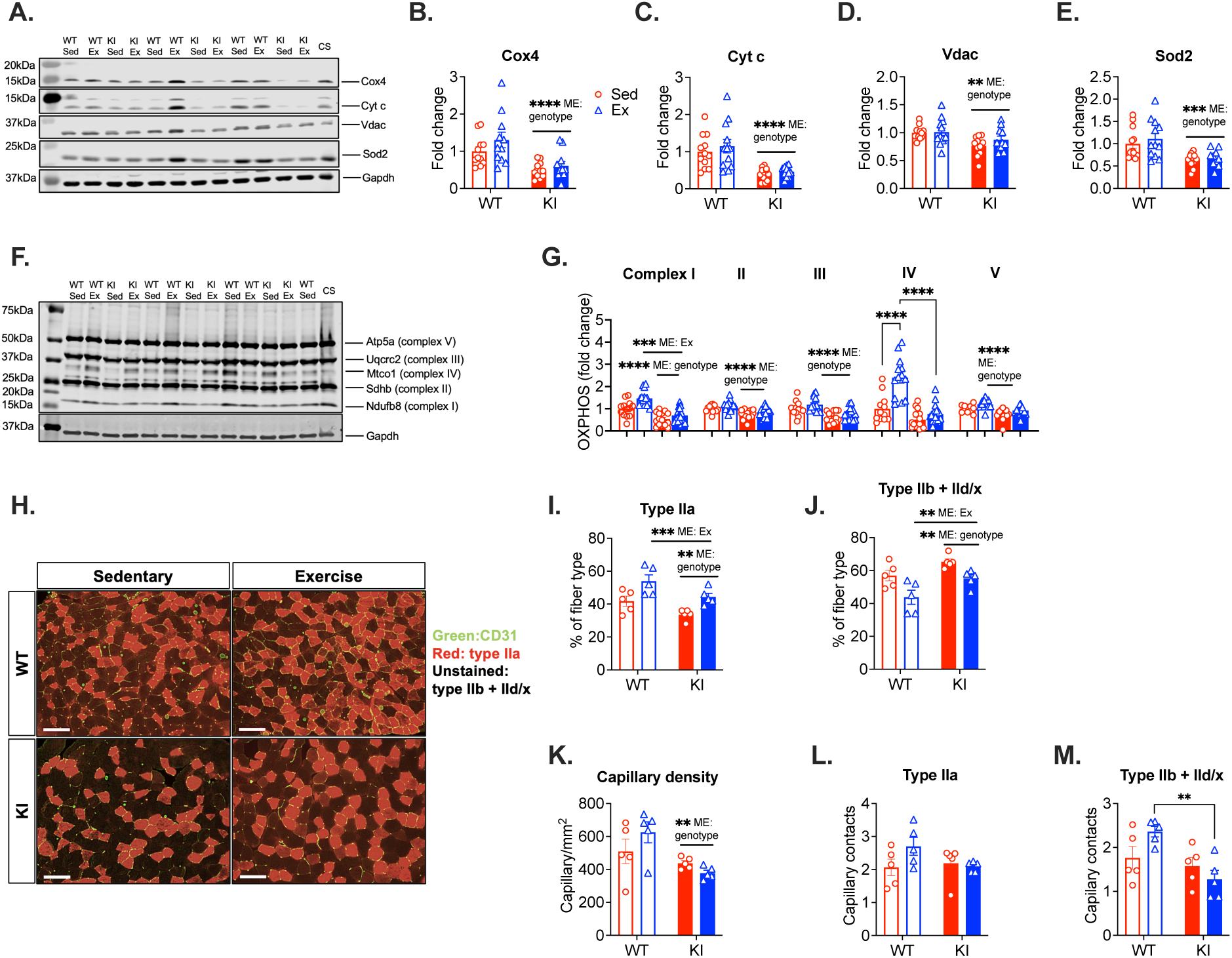
Mitochondrial and angiogenic adaptations, but not fiber type transformation, to exercise training is dependent on Ampkα2 T172 activation. Plantaris muscles from exercise-trained and sedentary mice were analyzed by western blot and immunofluorescence. **A**: Representative western blot image for Cox4, Cyt c, Vdac and Sod2; **B-E**: Quantification of protein expression in plantaris muscle; **F**: Representative western blot image for Oxphos complexes (I-V); **G**: Quantification of Oxphos subunit protein expression in plantaris muscle, n = 11-12 per group; **H**: Immunofluorescence staining of endothelial cells (green) and myosin heavy chain (MHC) type IIa in plantaris muscle (x20 magnification). Scale Bar = 100 μm; **I-J**: Quantification of percentage of type IIa (red) and type IIb + IId/x (unstained) fibers; **K-M**: Quantification of capillary density, capillary contacts per fiber for type IIa and type IIb + IId/x fibers, n = 5 per group. Data presented as means ± SEM and analyzed by two-way ANOVA. \*\**p* < 0.01, \*\*\**p* < 0.001, \*\*\*\**p* < 0.0001.

Next, we examined the muscle fiber-type composition in plantaris muscle. At baseline, sedentary KI mice exhibited a shift toward glycolytic fibers, characterized by increased type IIb and IId/x myofibers and diminished type IIa myofibers as compared to WT mice (Fig. 2*H-J*). VWR induced a similar increase in type IIa myofibers and reduction in type IIb and IId/x myofibers (Fig. 2*H-J*). These findings indicate that fiber-type transformation occurred independently of AMPKα2 T172 phosphorylation. Finally, we assessed angiogenesis. Exercise training increased capillary density (CD31) in WT mice; however, this effect was absent in KI mice (Fig 2*H* and *K*). A similar pattern was observed when analyzed across individual fiber types (Fig 2*L* and *M*). Collectively, these results demonstrate that AMPKα2 T172 phosphorylation is essential for mitochondrial and angiogenic adaptations to endurance training, but dispensable for fiber-type transformation in skeletal muscle.

### Endurance training-induced increase in skeletal muscle mitochondrial volume density is dependent on AMPKα2 T172 activation

We then utilized transmission electron microscope (TEM) to assess effects on mitochondrial reticulum morphology. VWR increased mitochondrial volume density (Mito_VD_) in plantaris muscle of WT mice but not in KI mice (Fig. 3*A* and *B*, S5*A*), whereas apparent mitochondrial number profile counting (Mito_P_) was unchanged across groups (Fig. 3*C*, S5*B*). Similar to the effect of endurance training on Mito_VD_, analyses of mitochondrial reticulum dimension, feret diameter and perimeter, showed trends for training-induced increases in WT but not in KI mice, with no changes observed in aspect ratio or circularity (Fig. 3*D-I*, S5*C-G*). Hence, these findings further support our concept that endurance training enhances muscle mitochondrial content, and this adaptation requires AMPKα2 T172 phosphorylation. Combined with mitochondrial protein data, these results support a conclusion that AMPKα2 activation plays a critical role in exercise induced mitochondrial remodeling, specifically mitochondrial biogenesis.

**Figure 3.**
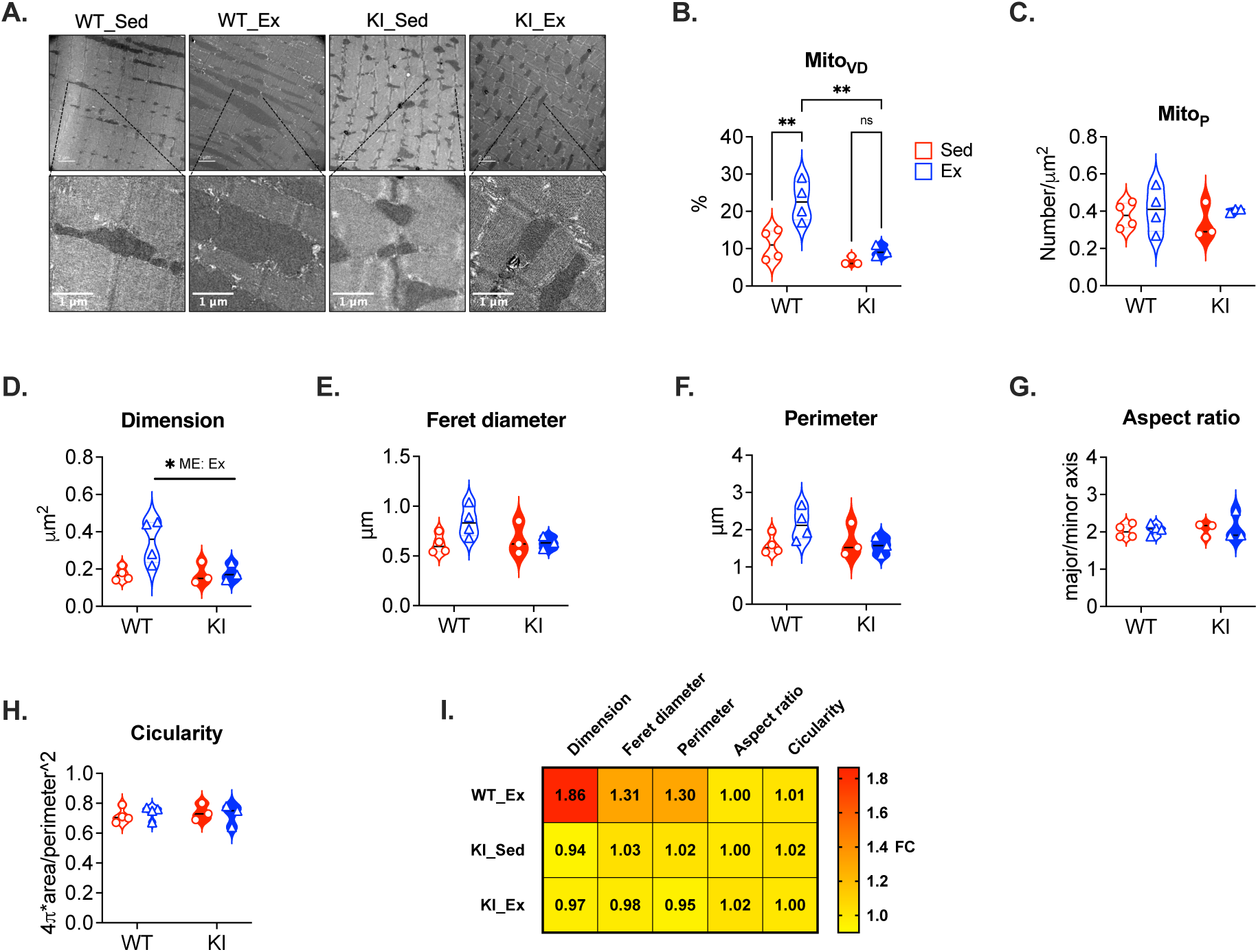
**Exercise training increases mitochondrial volume density in an Ampkα2 T172-dependent manner**. Intermyofibrillar mitochondria in the plantaris muscles from exercise-trained and sedentary mice were analyzed using transmission electron microscopy (TEM). **A**: Representative TEM images of WT and KI mice with or without exercise training. Scale bar = 1 μm; **B**: Mitochondrial volume density (Mito_VD_); **C**: Mitochondrial number profile counting; **D-H**: Additional measurements of mitochondrial morphology. A total of 350-500 mitochondria were analyzed from each mouse, and data are presented as the mean of all mitochondria from each mouse. **I**: A heatmap of quantification of mitochondrial morphology measurements normalized to sedentary WT mice, n = 3-4 per group. Data presented as means ± SEM and analyzed by two-way ANOVA. * *p* < 0.05, ** *p* < 0.01.

### AMPKα2 T172 activation is critical for exercise training-induced improvements in mitochondrial maximal respiratory function and metabolic flexibility in skeletal muscle

To investigate the role of AMPKα2 T172 activation in skeletal muscle mitochondrial bioenergetics and redox regulation following exercise training, we performed a high-resolution respirometry on mitochondria isolated from plantaris muscle using a creatine kinase clamp (Fig. 4*A*-*B*). Endurance training resulted in an increase of maximal respiration and enhanced respiratory conductance in WT mice, both of which were absent in KI mice (Fig. 4*C*-*F*). In contrast, KI mice exhibited significantly elevated mitochondrial H_2_O_2_ emission and electron leak across a range of energetic demands, indicating impaired redox balance (Fig. 4*G-H*). Finally, the impact of endurance training was minimal for redox regulation in both WT and KI mice. Mitochondrial ADP sensitivity, assessed by Michaelis-Menton kinetics for Km, was not altered by training or genotype (Fig. S6*A* and *B*). These findings further illustrate the importance of AMPKα2 T172 activation in exercise training-induced mitochondrial bioenergetic adaptation in skeletal muscle.

**Figure 4.**
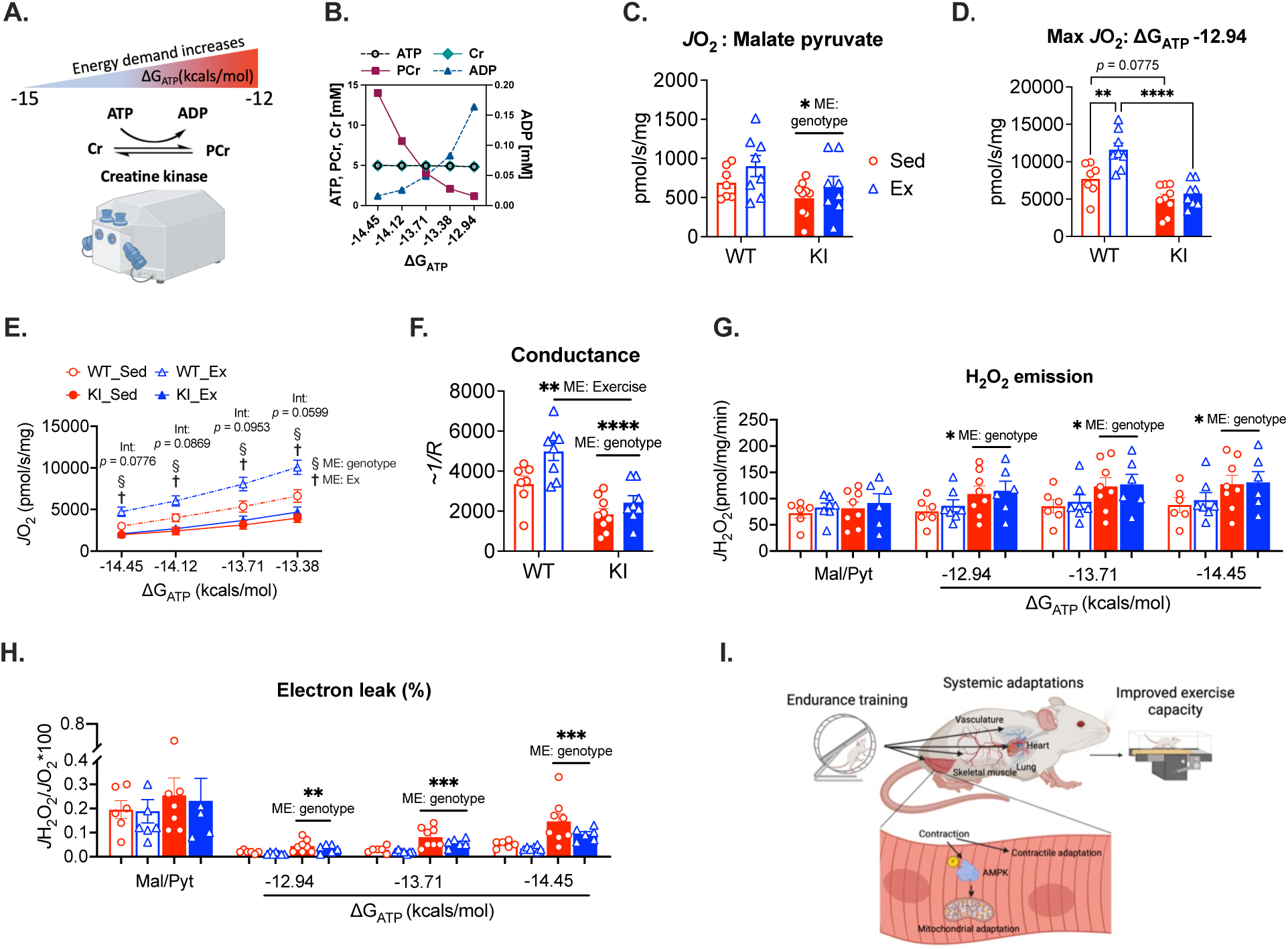
Exercise training improves mitochondrial respiratory function dependent on Ampkα2 T172 activation. Mitochondria isolated from plantaris muscle were assessed for respiratory function by high-resolution respirometry and reactive oxygen species (H_2_O_2_) emission by fluorospectroscopy in a creatine kinase clamp. **A-B**: A graphic illustration depicting the use of the creatine kinase clamp method, highlighting its key components and showing how the clamp modulates the concentrations of ATP, phosphocreatine (PCr), creatine (Cr) and ADP in response to varying bioenergetic demands. The illustration should demonstrate changes in ΔG_ATP_ (Gibbs free energy), which is highest at-12.95 and progressively declines (more negative) as energetic demand decreases; **C**: *J*O_2_ measurement at non-phosphorylating (NP) respiration; **D**: Maximal oxygen consumption (*J*O_2_ Max); **E**: Linear relationship between *J*O_2_ and ΔG_ATP_ during PCr titration; **F**: Conductance rate of PCr titration; **G**: H_2_O_2_ emission measurement; **H**: electron leak; n = 7-9 per group. Data presented as means ± SEM and analyzed by two-way ANOVA. \**p* < 0.05, \*\**p* < 0.01, \*\*\**p* < 0.001, \*\*\*\**p* < 0.0001; and **I**: A graphic demonstration summarizing the findings in the present study for potential factors determining exercise capacity following endurance training,

## Discussion

Repeated bouts of exercise activate multiple signaling networks in skeletal muscle to induce adaptive changes that improve functional performance (60, 61). Although AMPK has been shown to be a central driving factor in endurance exercise-induced mitochondrial and metabolic adaptations in skeletal muscle (34, 47, 51, 62–65), much of the evidence derives from mouse genetic models wherein protein stoichiometry was disrupted (44–47). Using a CRISPR-Cas9-mediated gene editing targeting the Ampkα2 T172 phosphorylation site, we previously demonstrated its essential role in overall energy metabolism at baseline and during exercise (51). In the present study, we further examined the role AMPKα2 T172 in exercise training-induced adaptations. Our data revealed that AMPKα2 T172 phosphorylation is required for exercise training-induced increases in mitochondrial content and respiratory function, as well as angiogenesis in skeletal muscle, but not for fiber type shifting or transformation. Despite impaired mitochondrial and angiogenic adaptations, *Ampkα2(T172A)* KI mice exhibited normal improvements in endurance exercise capacity and metabolic homeostasis with training. Thus, while these findings reveal the importance of AMPKα2 T172 activation, they also challenge the prevailing view that mitochondrial and angiogenic adaptations in skeletal muscle are required for enhanced exercise capacity.

There are various instances of experimental evidence supporting our findings regarding the role of AMPK in mitochondrial adaptation in response to exercise. First, acute treadmill running-induced mRNA expression of mitochondrial and oxidative genes was attenuated in muscle-specific *Ampkα1/2* knockout (KO) mice (44). Furthermore, long-term VWR promoted mitochondrial respiratory and Sod2 proteins in WT mice but not in muscle-specific *Ampkα2* kinase dead (KD) mice (37). Here, we used a new model system and obtained comprehensive evidence by expanding our measurements to mitochondrial ultra-structure and respiratory function as well as reactive oxygen species generation. Our findings support the notion that AMPKα2 T172 activation is required for endurance training-induced mitochondrial adaptation in skeletal muscle. However, studies using genetically modified mice, including skeletal muscle-specific knockout of Ampkα subunit(s) or overexpression of an inactive *Ampkα2* (DN), reported little impact of the loss of AMPK on exercise training-induced increases of mitochondrial proteins (44, 45, 47). Those discrepancies may reflect altered signaling landscape due to the disruptive nature of those genetic models. In this context, our KI model provides a distinct advantage by preserving the stoichiometry of AMPK while selectively abolishing canonical activation at AMPKα2 T172.

The evidence of the role AMPK in exercise training-induced angiogenesis is skeletal muscle is less abundant and definitive. In one study, acute treadmill running led to increased expression of *Vegf* mRNA, an important regulator for capillary maintenance and angiogenesis, in an AMPKα-dependent manner (44), whereas, in another study, long-term voluntary wheel running resulted in increased capillary density and fiber-type transformation toward a more oxidative phenotype in skeletal muscle in both WT and muscle-specific transgenic mice of inactive *Ampkα2* (DN) with a greater induction of *Vegf* mRNA in DN mice compared with WT mice (66). Our findings are consistent with those of the first study. In a recent study we used phosphoproteomic analysis in skeletal muscle to reveal that *Ampkα2(T172A)* KI mice had a significant perturbation in eNOS (Nos3) signaling in response to an acute bout of exhaustive exercise (51). In fact, eNOS has been shown to be essential for ischemia-induced vascular remodeling in skeletal muscle (67), and inhibition of NOS activity blocked VEGF and VEGFR2 expression and angiogenesis induced by chronic electrical stimulation of motor nerve, an in vivo model mimicking endurance exercise (68). Our findings suggest that AMPKα2 T172 activation is required for endurance training-induced angiogenesis in skeletal muscle.

Considering present and previous results, a remaining question concerns the mechanism by which AMPK activation may trigger adaptive processes in skeletal muscles following each bout of endurance exercise. AMPK has been shown to directly phosphorylate PGC-1α (38), a master regulator of mitochondrial biogenesis and angiogenesis in skeletal muscle (69–71), and PGC-1α’s nuclear/mitochondrial translocation and subsequent transcriptional activities have been shown to be dependent on AMPK (72–74). Importantly, exercise induced-mitochondrial biogenesis and angiogenesis, but not fiber-type transformation, in skeletal muscle have been shown to be dependent on Pgc-1α (39, 40). It is important to note that our findings in the current study phenocopied these previous studies, supporting that AMPK activation in concert with PGC-1α in inducing mitochondrial biogenesis and angiogenesis, but not to be responsible for fiber type transformation in skeletal muscle in response to endurance training.

Increased mitochondrial content and respiratory enzyme activity and improved capillary density in skeletal muscle are classically linked to functional improvements by exercise training (2, 8, 14, 17, 19, 75). These adaptations may reduce metabolic perturbation during exercise with enhanced fat oxidation and lactate clearance, leading to an improved intramyocellular homeostasis and, thereby, endurance exercise capacity (12, 18). Consistently, a recent study by García-Domínguez et al. reported that mitochondrial adaptations are functionally required for exercise-induced improvements in exercise capacity (14). An intriguing and unexpected finding of the present study is that exercise training-induced mitochondrial and angiogenic adaptations were ameliorated in KI mice, yet endurance exercise capacity and substrate selection during VO_2_max testing were improved to a similar extent as in WT mice, which are very similar to the findings observed in mice with muscle-specific deletion of the master gene of mitochondrial remodeling*, Pgc-1α* (76). Thus, several interpretations of the data are possible. Our findings can be interpreted to mean that exercise-induced mitochondrial biogenesis and angiogenesis in skeletal muscle are not required for the improved maximal exercise capacity following endurance training in adult mice (Fig. 4*I*). It could also mean that respiratory capacity in mouse muscle is in surplus to begin with. And, finally, unmeasured factors, such as sarcolemmal and mitochondrial monocarboxylate transporter (MCT) abundances (77) that are involved in lactate clearance (53, 54, 78), were involved.

Over the years there have been suggestions that central factors, such as cardiac output, pulmonary system and blood flow, are main determining factor for 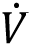O_2_max and endurance capacity in healthy, untrained adult subjects (13, 79, 80), whereas peripheral factors, such as mitochondrial mass in skeletal muscle, may become limiting during endurance exercise, aging and disease states (81, 82). Alternatively, classic and current findings may be interpreted to mean that our bodies are elegantly built with multiple redundancies in the regulation of carbon and oxygen flows, and their matching, under diverse circumstances. Thus, as in the present investigation, alteration of a single gene may produce ambiguous results when obviously multiple enzyme and organ systems are involved in controlling exercise endurance under diverse circumstances

## Conclusion

Phosphorylation of AMPKα2 T172 is a critical regulatory step in the structural and bioenergetic remodeling processes in skeletal muscle in response to endurance training in young, healthy animals. However, these adaptations are not obligatorily required for the improvement of exercise capacity following training. These findings reinforce the complexity and elegant redundancy of whole-body functional adaptation to exercise training.

## Materials and Methods

### Animals

All experimental procedures were approved by the Institutional Animal Care and Use Committee at the Virginia Tech (protocol No. 22-251). To ascertain the functional role of the Ampkα2 (threonine 172), we performed CRISPR/Cas9-mediated gene editing and developed loss-of-function *Ampkα2 (T172A)* mice with mutation of the critical activation phosphorylation site threonine to alanine. For experiments in the present study, *Ampkα2 (T172A)* KI mice (in C57BL/6 background) were generated at the Genetically Engineered Murine Model (GEMM) core facility at the University of Virginia. Both sexes of *Ampkα2 (T172A)* KI and age-matched WT C57BL/6 controls were used for experiments. Mice were assigned randomly to different experimental groups (n = 17-18; male = 9-10/group, female = 7-9/group). Sedentary mice were group housed (2-4 mice per cage), exercise trained mice were housed individually in VWR cages, in temperature-controlled (21 °C) quarters with a 12:12-h light-dark cycle and ad libitum access to water and normal chow (Purina).

### Acute treadmill running

Acute endurance exercise was conducted by using the Exercise 3/6 Treadmill (Columbus Instruments, Columbus, OH, USA) following a protocol described before (51, 83).

### Electrical stimulation

Nerve-stimulated contraction was conducted in vivo under anesthesia using the Aurora Whole Animal System according to our previously published procedure (84).The frequency stimulation was consisted of a 10 Hz stimulation frequency, 0.25 ms stimulation duration for 30 min per session for 4 sessions (2 hrs in total). Immediately after the electrical stimulation procedure, plantaris muscle tissue was collected and stored at-80°C upon further analysis.

### Exhaustive treadmill running test

Endurance capacity was assessed by an exhaustive treadmill running test for all experimental mice according to our previous published studies (83, 85, 86).

### Glucose tolerance test

Glucose tolerance test was performed according to our published procedure (83) 24 hrs after the last bout of voluntary wheel running.

### VO2max treadmill running test

All mice were tested by the Oxymax Metabolic Treadmill System (Columbus Instruments, Columbus, OH, USA) (51).

### Tissue preparation & Western Blot

Mice were euthanized under 2.5% isoflurane at the time of skeletal muscle harvest. Mice were fasted overnight before tissue collections. The plantaris, gastrocnemius, soleus, heart and tibia were removed with surgical tools rinsed in 1X PBS, weighed and prepared for different assays accordingly. For western blot, tissues were homogenized in glass homogenizers with protein sample buffer containing 50 mM Tris-HCL, pH 7.4, 0.01% bromophenol blue, 10% glycerol, 1% sodium dodecyl sulfate (SDS), 127 mM 2-mercaptoethanol, and 20 mM dithiothreitol, supplemented with protease inhibitor cocktails and phosphatase inhibitor cocktail tablets (Sigma-Aldrich). Homogenized tissue lysates were then boiled in a heat block at 98 °C for 5 min and stored in-70 °C. Protein concentration was determined by the RCDC protein assay kit (Biorad). Protein lysates were subjected to sodium dodecyl sulfate-polyacrylamide gel electrophoresis at 100V for ∼2 hrs, and transferred onto nitrocellulose membranes at 80V for 2 hrs. Membranes were probed with the following primary antibodies at a 1:1000 dilution: targeting phospho-Ampkα1/2 (Thr172) (CST, 2535), Ampkα1/2 (CST, 2532), phospho-Acc (CST, 3661), phospho-Ulk1(CST, 5869), Ulk1 (Sigma-Aldrich, A7481), phospho-Akt (CST, 4060), Akt (CST, 9272), Cox4 (CST, 4844), cytochrome c (CST, 4272), Vdac (CST, 4866), Sod2 (CST, 13141), total Oxphos cocktail (Abcam, ab110413) Apool (Proteintech, 28514-1-AP), Chchd3 (Proteintech, 25625-1-AP), Immt (Proteintech, 10179-1-AP), Samm50 (Proteintech, 28679-1-AP), Gapdh (CST, 2118). Secondary antibodies were goat anti-rabbit IR800 and anti-mouse IR680. Membranes were scanned using the odyssey infrared imaging system (LICOR). Proteins were analyzed and normalized to a common protein standard loaded on gel.

### Immunofluorescence

Skeletal muscle fiber type and capillary density were determined by immunofluorescence staining in plantaris muscles as previously described (17, 39). Rat anti-CD31 (MCA1364, Serotec) followed by mouse anti-MHC I (BF-F8) and anti-MHC IIa antibody (SC71), followed by appropriate fluorophore-conjugated secondary antibodies. Images were acquired using a Keyence BZ-X800 Series Fluorescence Microscope under identical settings across samples. Muscle fiber types were classified based on fluorescence signals, with type I and type IIa fibers identified by specific staining, and unstained fibers categorized as type IIb + IId/x. Fiber type distribution was quantified across the entire muscle cross-section at x4 magnification. Capillary density was assessed by quantifying CD31-positive puncta relative to tissue area at 20x magnification. In addition, capillary contacts per fiber type were determined by counting the number of capillaries surrounding individual fiber types.

### Transmission electron microscope (TEM)

Plantaris muscle tissue were harvested from mice and prepared accordingly to TEM core facility. Muscle tissues were fixed overnight at 4 °C in 2.5% glutaraldehyde and 4% paraformaldehyde in 0.1 M NaCacodylate buffer (pH 7.4). After buffer washes, samples were post-fixed in 2% osmium tetroxide with 1.5% potassium ferrocyanide for 1h at room temperature, followed by 1% uranyl acetate stain in 50% ethanol for 1 h. Dehydration was carried out in graded ethanol (70%, 90%, 100%), then tissue were infiltrated with the EPON/propylene oxide mixture overnight and followed by embedded in 100% EPON resin. Ultrathin sections were cut and collected on copper grids. Samples were examined under the FEI Tecnai (G2 Spirit Twin) transmission electron microscope (ThermoFisher Scientific, Hillsboro, OR, USA) at 120 kV with a magnification of x1200. To avoid sampling bias, all samples were assigned anonymous codes, and all analyses were conducted blindly. Mitochondrial features were quantified using a two-dimensional serological method. For each sample, ten random images were acquired and analyzed with NIH imageJ software. A defined 9×9 grid was superimposed on each image. The relative mitochondrial volume density (Mito_VD_), define as the volume of mitochondria per volume of muscle fiber, was estimated by counting the number of grid intersections falling on mitochondria and dividing by the total number of grid points. Mitochondrial number (mitochondrial profiles per area, Mito_P_) was quantified by the number of mitochondria per unit cytoplasmic area. Individual mitochondrial features were quantified by free hand-drawing tool of ImageJ to quantify mitochondrial reticulum dimension, feret diameter, perimeter, aspect ratio and circularity.

### Preparation of Isolated mitochondria from skeletal muscle

The procedure of isolating mitochondria was described in our previous published study (87). Briefly, mice plantaris muscle were harvested and then, homogenized in a 1000 mL of cold FRAC buffer (0.03 mM fatty acid-free BSA, 70 mM sucrose, 210 mM Mannitol, 5 mM HEPES, and 1 mM EGTA) through an electrical tissue homogenizer (BioSpec) at 4 °C for ∼30 seconds. Homogenized tissues were then spun at 800X g for 10 minutes at 4 °C. Supernatant were then collected and spun at 9000X g for 10 minutes at 4 °C. The resultant pellets were collected and resuspended in FRAC buffer (without BSA) for protein concentration determination by the Bradford assay (Thermo-Fischer, Agawam, MA, USA).

### High-resolution respirometry

High resolution respirometry was used to assess mitochondrial respiratory rate (*J*O_2_) or oxygen consumption rate (OCR) through Oroboros O2k (Innsbruck, Austria). The creatine kinase (CK) clamp technique was employed to evaluate *J*O_2_ of isolated mitochondria from plantaris muscle. Compared to conventional ADP derived approach, the CK technique takes advantage of enzymatic reaction of CK to clamp ATP:ADP under a physiologically relevant range (Fig. 4*A* and *B*)(88, 89). Through this platform, we were able to also measure the metabolic flexibility or mitochondrial conductance by having sequential phosphocreatine (PCr) titrations to create different levels of ATP free energies (ΔG_ATP_). Theoretically, *J*O_2_decreases when ΔG_ATP_ decreases and vice versa, when electron transport chain (ETC) is tightly coupled to the changing demand for ATP re-synthesis. On days of the experiment, Oroboros O2k chambers were warmed to 37°C and set to 500 rpm stirrer speed and filled with Buffer D (105 mM K-MES potassium salt, 30 mM KCl, 1 mM EGTA, 10 mM KH_2_PO_4_, 5 mM MgCl_2_-6H_2_O, 0.05% BSA, pH7.1) to record the 100% air saturation, followed by saturating sodium dithionite (Sigma S1256) to record zero-point calibration. Following calibration, chambers were washed thoroughly with water, then refilled with Buffer D supplemented with 5mM creatine monohydrate (Sigma C0780). 10ug of mitochondria based on the Braford assay of each sample was added into each chamber, energized by 5mM pyruvate and 2.5mM malate (non-phosphorylating respiration). Maximal respiratory flux was measured by adding 20mM creatine kinase, 1mM PCr and 5mM ATP, which generates a ΔG_ATP_ of-12.94 kcal/mol. Cytochrome C (0.005mM) was added to evaluate the membrane integrity of mitochondria with a threshold of 10% increase in flux to be exclusionary. Finally, sequential PCr titrations was performed (3, 6, 15, 30mM) to create 4 different free energy states (ΔG_ATP_ =-13.38,-13.71,-14.12, - 14.45 kcal/mol, respectively) to evaluate the corresponding mitochondrial respiratory flux changes (conductance). The calculation of ΔG_ATP_ for CK bioenergetic clamp can be found here: https://dmpio.github.io/bioenergetic-calculators/ck_clamp/.

We evaluated mitochondrial ADP sensitivity defined by [ADP]_50_ Km, the ADP concentration required to reach half maximal respiratory flux, by using Michaelis-Menten kinetics. ADP concentrations were calculated as previously described (90).

### Measurement of mitochondrial ROS emission

Mitochondrial reactive oxygen species (ROS) emission was determined using Amplex Ultra Red (AUR), based on the concept that horseradish peroxidase (HRP) catalyzes the Hydrogen peroxide (H_2_O_2_)-dependent oxidation of non-fluorescent Amplex Red to fluorescent resorufin. Superoxide dismutase was added to the reaction mixture to ensure that all superoxide is converted to H_2_O_2_. Briefly, a standard curve for H_2_O_2_ (25-250 nM) was prepared using buffer D, containing AUR stock solution (0.02 mM; prepared according to the manufacturer’s instructions), HRP (1U/mL), SOD (20 U/mL), auranofin (AF; 0.0001 mM), malate (2.5 mM), pyruvate (5 mM), creatine kinase (20 mM), phosphocreatine (PCr; 1mM), and ATP (5 mM) in a final volume of 0.6 mL. Fluorescence measurements were performed using a Horiba Quantmaster 4 Fluorometer (Edison, NJ, USA) under the following conditions: 37 °C, excitation/emission wavelengths of 565/600 nm, stirring at 500 rpm, and a 0.875 mL cuvette with an adaptor. Data collection set to 1-second intervals with a 60-second pause, over a 4-minute period for each stage. The H_2_O_2_ concentration in the reaction will range from 25 to 250 nM, using a 0.1 mM H_2_O_2_ stock solution. The background fluorescence was subtracted using FelixFL software after 10 seconds of exposure prior to AUR addition with an open window. Further background correction was applied to each run based on the fluorescence rate before H_2_O_2_ addition. The H_2_O_2_ concentration was calculated as previously described range (88, 89). Following mitochondrial isolation from the plantaris and protein quantification (described above), 10 μg of mitochondria each sample was added to the pre-warmed cuvette under the same conditions as the O2k assay. Electron leak was expressed as a percentage of total flux, calculated as the ratio of H_2_O_2_ production (*J*H_2_O_2_) to oxygen consumption (*J*O2) (*J*H_2_O_2_ / *J*O2 = % leak).

## Statistical Analyses

Data are presented as means ± SE. Experimental outcome measures with only one variable was analyzed using Student’s *t* test. When there are two variables (Exercise × Genotype), outcome measures were analyzed via two-way ANOVA, with Tukey post hoc analyses when there is a significant interaction. Statistical significance was set as *p* < 0.05.

## Acknowledgments

We thank Wenqing Shen, A Robb Burgart and Mei Zhang for their technical support.

## Author Contributions

X.M. and Z.Y. conceived and designed the studies; X.M. and R.N.M. performed the experiments; X.M., R.N.M., and K.T. analyzed the data; X.M., R.N.M., and Z.Y. interpreted the results of experiments; X.M., R.N.M. and Z.Y. prepared the figures; X.M. drafted the manuscript; X.M., R.N.M., K.T., F.W.B., G.A.B. and Y.Z. edited and revised the manuscript; X.M., R.N.M., K.T., F.W.B, G.A.B. and Z.Y. approved final version of the manuscript.

## Data sharing plans

All data is included in the manuscript and/or supporting information.

## Supplementary information/figures

**Supplementary figure 1.**
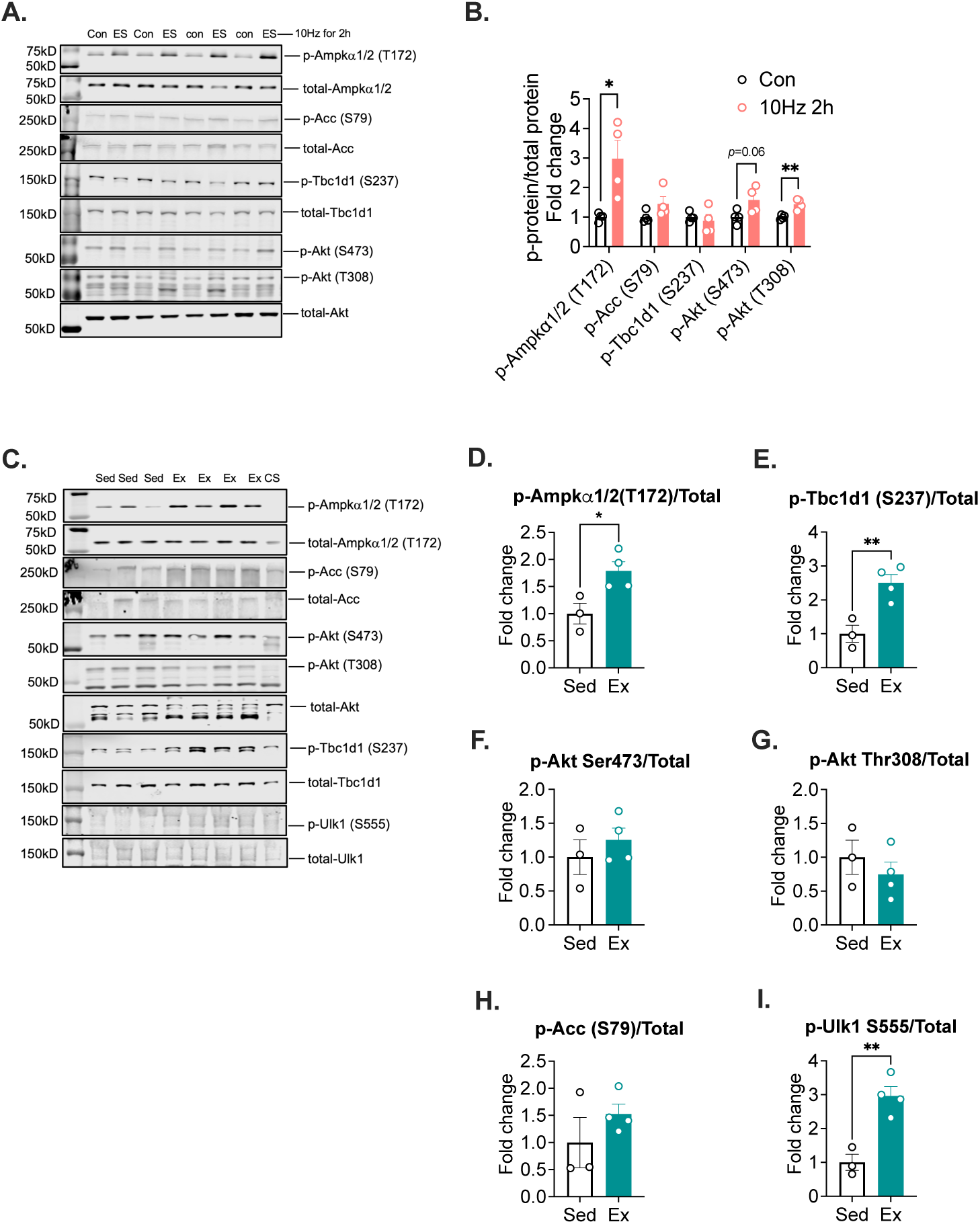
Acute treadmill running and electrical stimulation activate Ampkα T172 in skeletal muscle. Wild type C57BL/6J mice (male) were subjected to 90 min acute treadmill running with sedentary mice as control or unilateral electrical stimulation of the sciatic nerve at 10 Hz for 2 hours with the contralateral leg as control. Plantaris muscles were analyzed by western blot. **A**: Representative western blot images for p-Ampkα1/2 (T172), p-Acc (S79), p-Tbc1d1 (S237), p-Akt (S473 & T308), and respective total proteins in electrically stimulated muscle (ES) and the contralateral control muscle (Con); **B**: Quantification of protein expression, n = 4; **C**: Representative western blot images for p-Ampkα1/2 (T172), p-Acc (S79), p-Akt (S473 & T308), p-Tbc1d1 (S237), p-Ulk1 (S555), and respective total proteins from exercised (Ex) and sedentary mice (Sed), CS stands for common standard; **D-I**: Quantification of protein expression, n = 3-4 per group. Data presented as means ± SEM and analyzed by two-tailed *t*-test. \**p* < 0.05, \*\**p* < 0.01.

**Supplementary figure 2.**
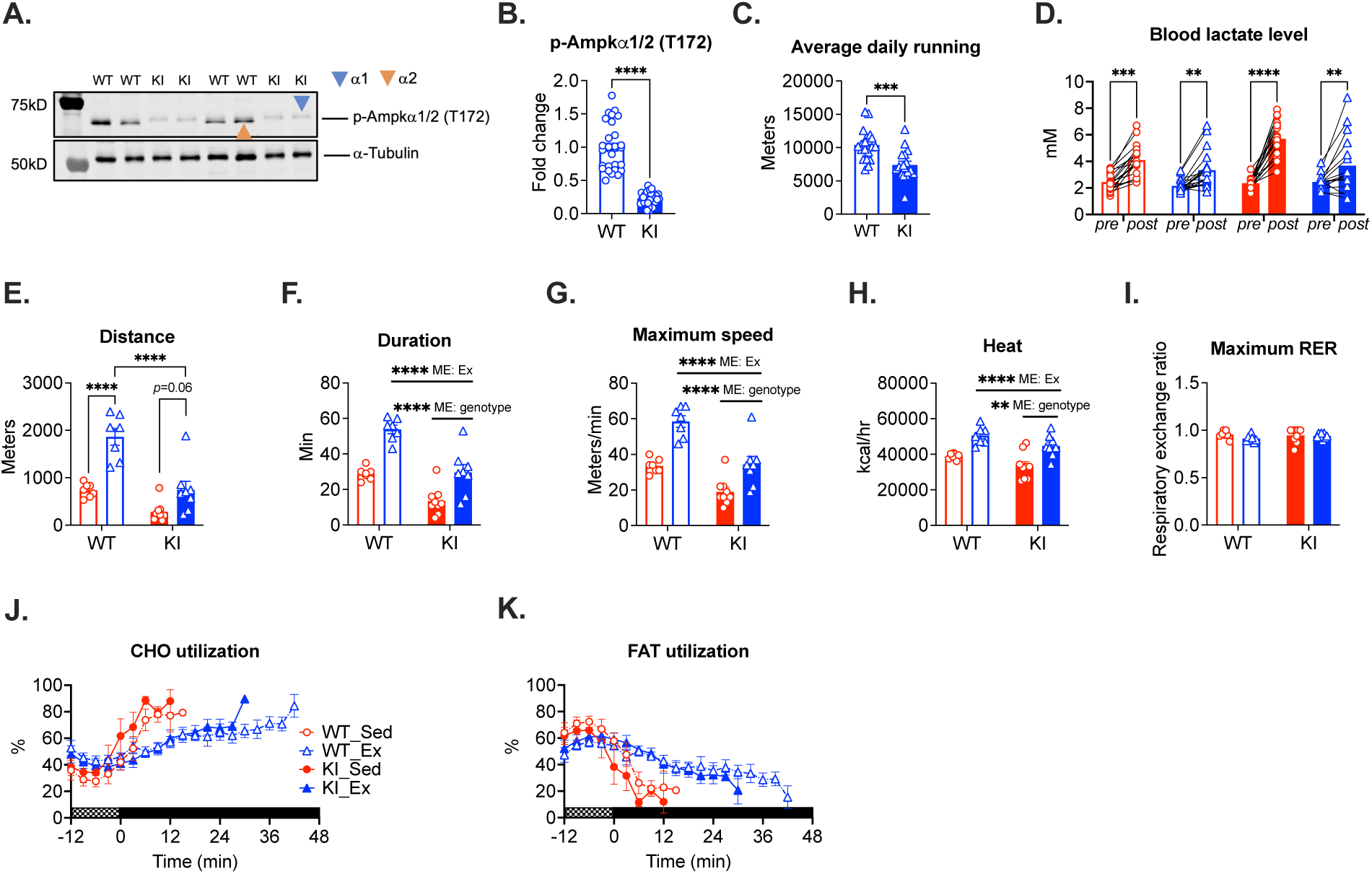
Additional parameters measured during exhaustive treadmill running test and VO2max treadmill running test. *Ampkα2*(T172A) KI and WT littermates were subjected to VWR for 4 weeks with sedentary KI and WT mice as controls followed by exhaustive treadmill running test and VO_2_max treadmill running test. **A**: Representative western blot image for p-Ampkα1/2 (T172) with α-tubulin as loading control in plantaris muscles. Blue triangle points to α1 subunit, and orange triangle points to α2 subunit; **B**: Quantification of the p-Ampkα1/2 (T172) in WT and *Ampkα2*(T172) KI mice; **C**: Average daily voluntary wheel running distance for WT and KI mice, n = 18-20 per group. Data presented as means ± SEM and analzyed by two-tailed *t*-test. *** *p* < 0.001, **** *p* < 0.0001; **D**: Blood lactate before and after exhaustive treadmill running test; **E-I**: Total running distance, duration, maximal speed, heat production, and maximum RER during the VO_2_max treadmill running test; and **J-K**: Percentage of carbohydrate and fatty acid oxidation, n = 7-9 per group. Data presented as means ± SEM and analyzed by two-way ANOVA. ** *p* < 0.01, **** *p* < 0.0001.

**Supplementary figure 3.**
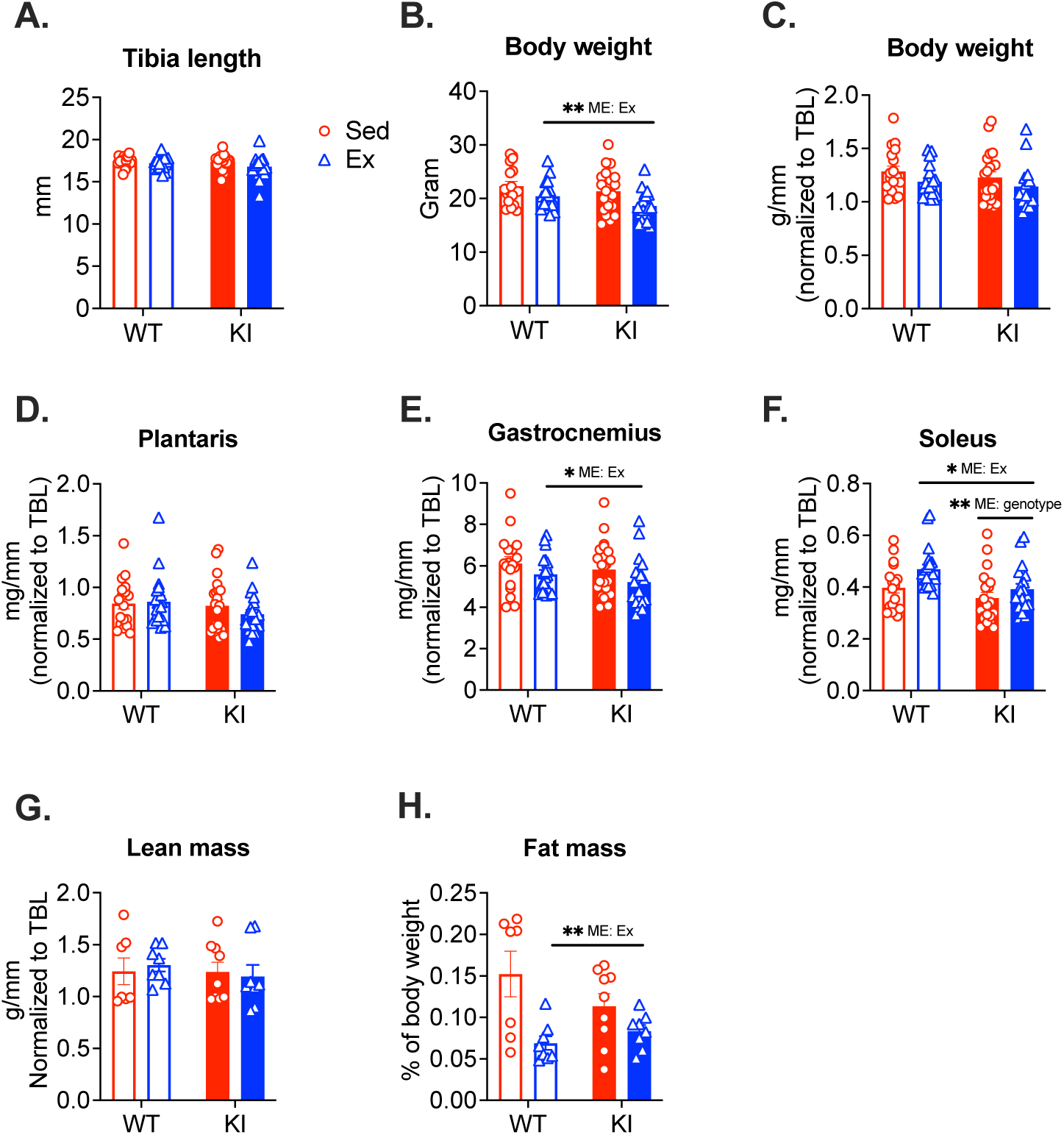
Measurements of body weight, muscle mass, lean body mass and fat mass in exercise trained and sedentary WT and *Ampkα2*(T172) KI mice. *Ampkα2*(T172A) KI and WT littermates were subjected to VWR for 4 weeks with sedentary KI and WT mice as controls followed by direct measurements of tibia length, body weight and muscle mass and echoMRI measurments of lean body mass and fat mass. **A**: Tibia length (TBL); **B-C**: Body weight and body weight normalized by TBL; **D-F**: Skeletal muscle mass nomalized by TBL, n = 18-20 per group; **G-H**: Lean body mass and fat mass, n = 7-9 per group. Data presented as means ± SEM and analyzed by two-way ANOVA. * *p* < 0.05, ** *p* < 0.01.

**Supplementary figure 4.**
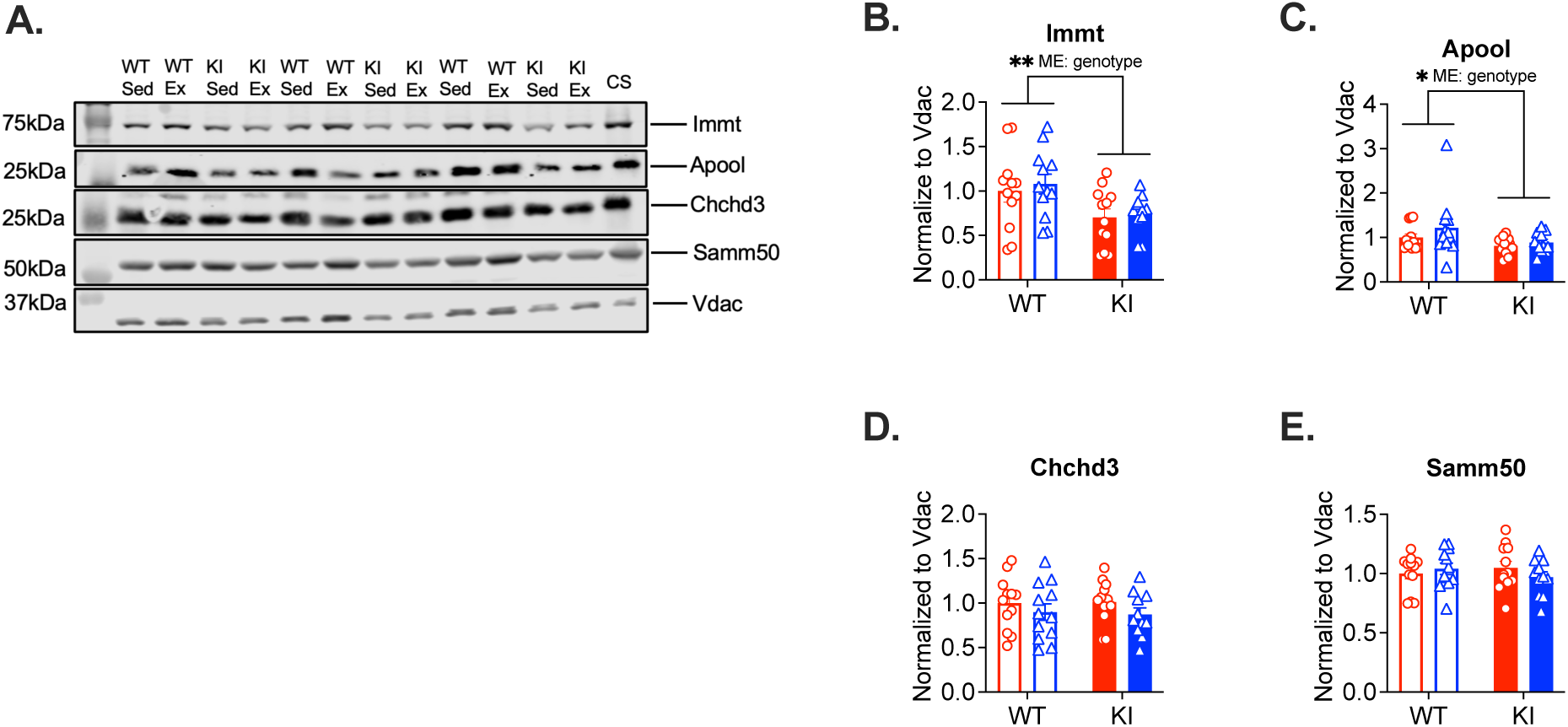
Mitochondrial contact site and cristae organizing system (MICOS) proteins are reduced in *Ampkα2*(T172A) mice. *Ampkα2*(T172A) KI and WT littermates were subjected to VWR for 4 weeks with sedentary KI and WT mice as controls followed by western blot analysis of plantaris muscle. **A**: Representative western blot image for Immt (Mic60), Apool (Mic27), Chchd3 (Mic19) and Samm50; **B-E**: Quantification of protein expression normalized by Vdac, n = 11-12 per group. Data presented as means ± SEM and analyzed by two-way ANOVA are * *p* < 0.05, ** *p* < 0.01.

**Supplementary figure 5.**
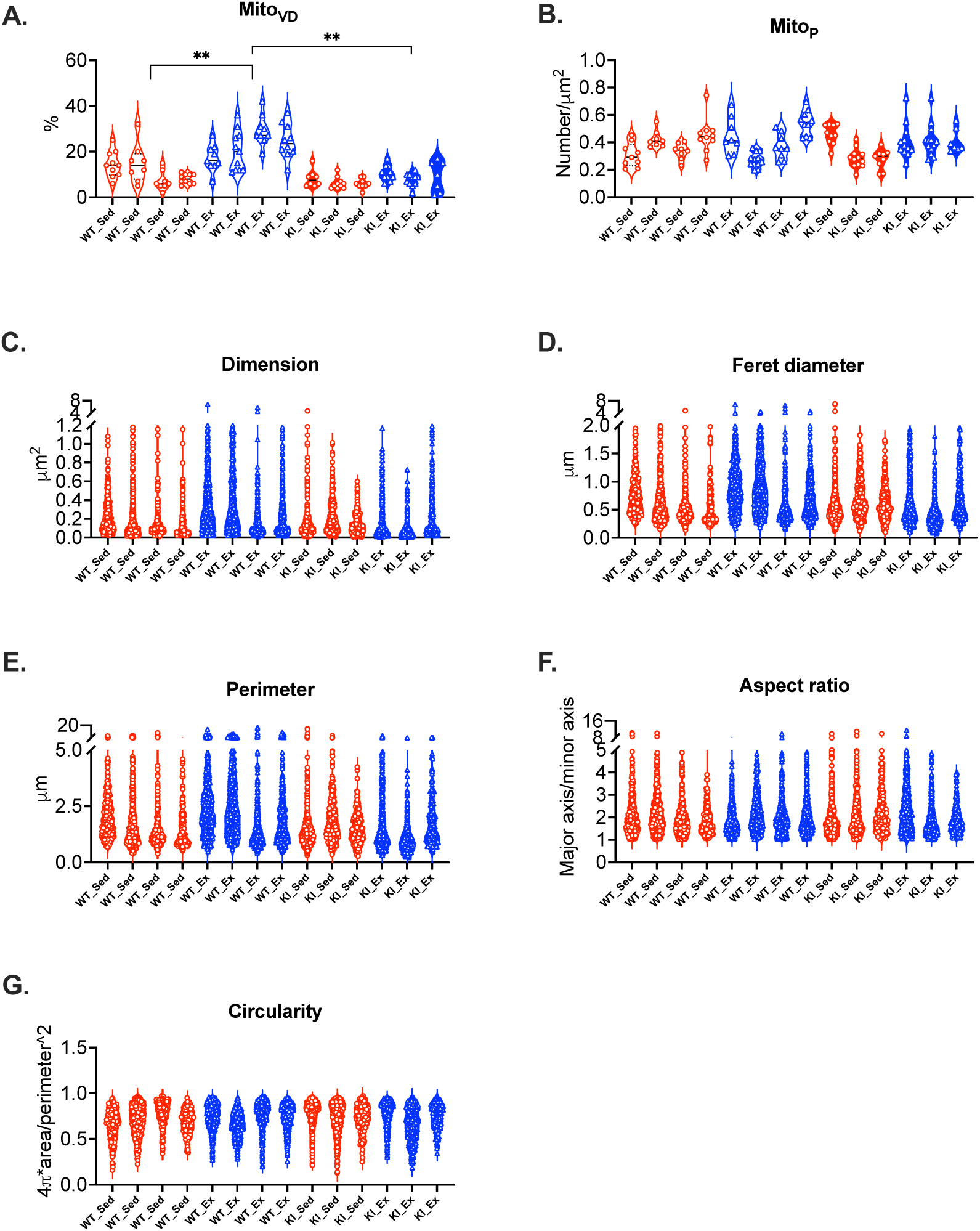
Measurements of mitochondrial morphology and structure by TEM in plantaris muscle. *Ampkα2*(T172A) KI and WT littermates were subjected to VWR for 4 weeks with sedentary KI and WT mice as controls followed by TEM analysis in plantaris muscles. **A-B:** Quantification of mitochondrial volume density (Mito_VD_) and number (Mito_P_). Each dot denotes a single image analyzed, and each column denotes a single mouse; **C-G**: Mitochondrial morphological measurements (dimension, feret diameter, perimeter, aspect ratio and circularity) from each mouse. Each dot denotes one mitochondrion (350-550 mitochondria quantified per mouse). Data presented as means ± SEM and analyzed by two-way ANOVA. ** *p* < 0.01.

**Supplementary figure 6.**
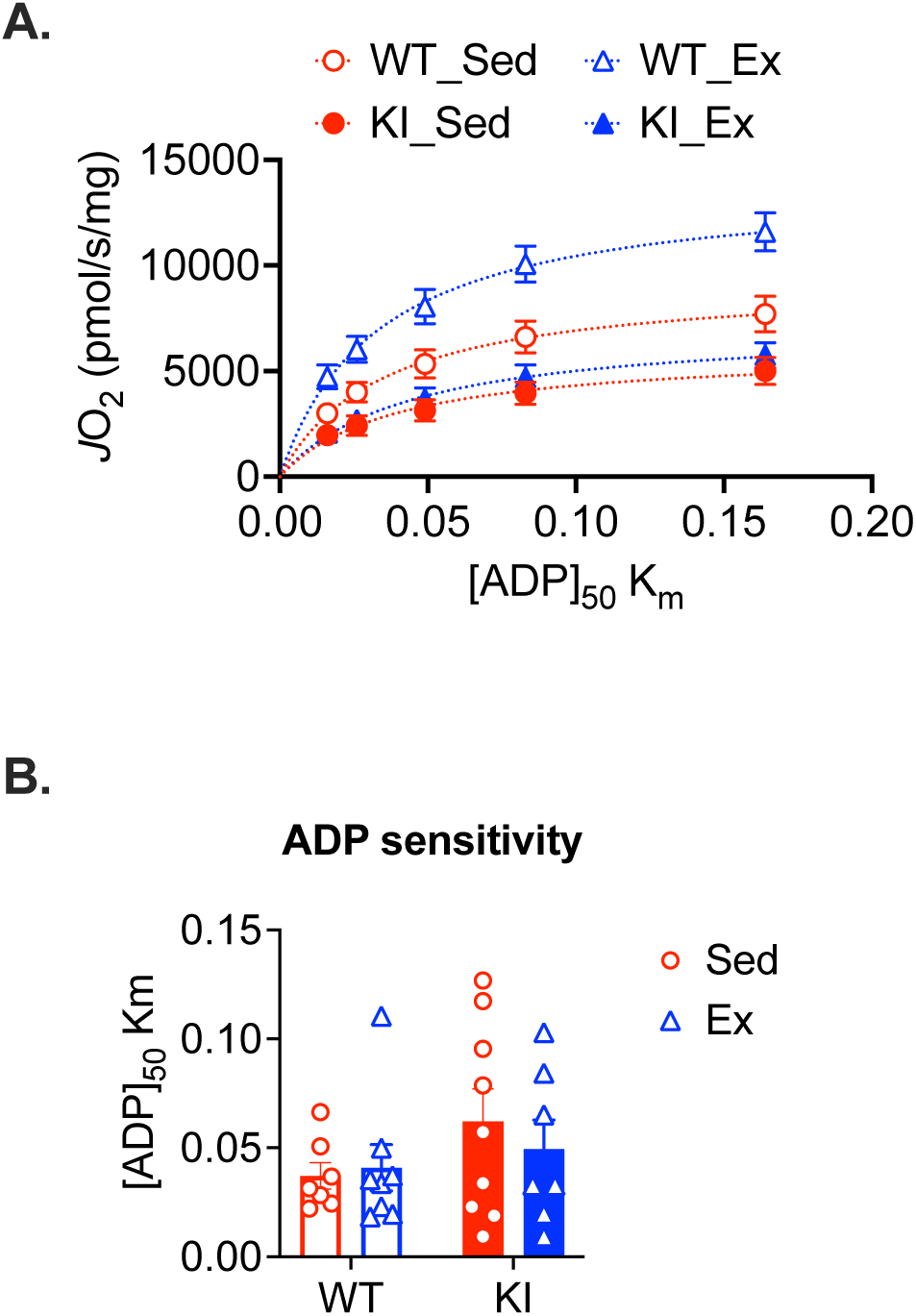
ADP sensitivity is not significantly changed by exercise or Ampkα2 (T172). *Ampkα2*(T172A) KI and WT littermates were subjected to VWR for 4 weeks with sedentary KI and WT mice as controls followed by high-resolution respirometry analysis of isolated mitochondria from plantaris muscles. **A**: Oxygen consumption at ADP [c] 0.016mM, 0.026mM, 0.049mM, 0.083mM, and 0.164mM corresponding to ΔG_ATP_ of-14.45,-14.12,-13.71,-13.38, and-12.94kCal/mol, respectively; and **B**: ADP sensitivity (Km) determined by Michaelis Menton analysis, n = 7-9 per group. Data presented as means ± SEM.

